# Loss of neuronal Imp induces seizure behavior through Syndecan function

**DOI:** 10.1101/2024.11.21.624719

**Authors:** Paula R. Roy, Nichole Link

**Affiliations:** Department of Neurobiology, University of Utah, Salt Lake City, UT, 84112, USA

## Abstract

Seizures affect a large proportion of the global population and occur due to abnormal neuronal activity in the brain. Unfortunately, widespread genetic and phenotypic heterogeneity contribute to insufficient treatment options. It is critical to identify the genetic underpinnings of how seizures occur to better understand seizure disorders and improve therapeutic development. We used the *Drosophila* model to identify that IGF-II mRNA Binding Protein (Imp) is linked to the onset of this phenotype. Specific reduction of Imp in neurons causes seizures after mechanical stimulation. Importantly, gross motor behavior is unaffected, showing Imp loss does not affect general neuronal activity. Developmental loss of Imp is sufficient to cause seizures in adults, thus Imp-modulated neuron development affects mature neuronal function. Since Imp is an RNA-binding protein, we sought to identify the mRNA target that Imp regulates in neurons to ensure proper neuronal activity after mechanical stress. We find that Imp protein binds *Syndecan* (*Sdc*) mRNA, and reduction of Sdc also causes mechanically-induced seizures. Expression of Sdc in *Imp* deficient neurons rescues seizure defects, showing that Sdc is sufficient to restore normal behavior after mechanical stress. We suggest that Imp protein binds *Sdc* mRNA in neurons, and this functional interaction is important for normal neuronal biology and animal behavior in a mechanically-induced seizure model. Since Imp and Sdc are conserved, our work highlights a neuronal specific pathway that might contribute to seizure disorder when mutated in humans.

## INTRODUCTION

Seizure disorders are debilitating but common conditions. An estimated ten percent of the human population will have at least one seizure in their life, and up to three percent of the global population experiences recurring spontaneous seizures defined as epilepsy^1^. Despite their prevalence, phenotypic heterogeneity makes it challenging to identify seizure causality. Therefore, the development of patient-specific treatments can be difficult, but an understanding of the genetic pathways underlying seizure pathology may aid in the development of patient directed therapies. Model organism research can greatly facilitate the identification of genetic-based seizure disorders^2,3^. Genetic pathways can be studied in detail, mechanisms of pathogenesis are often uncovered, and drugs can be screened in a patient-specific manner to identify candidate therapeutics.

A class of genes with growing connection to syndromes involving seizures are RNA-binding proteins^4^. Therefore, the contribution of RNA-binding proteins in neurodevelopment and neurological disease is an emerging interest in the fields of genetics and neuroscience. RNA-binding proteins spatially and temporally regulate gene expression to coordinate and modify dynamic cellular networks like those found in the brain^5,6^. Insulin Growth Factor 2 Binding Proteins (IGF2BPs) are mRNA-binding proteins that post-transcriptionally regulate mRNA by influencing localization, transport, translation, and stability^7,8^. Vertebrates have three IGF2BP paralogs (1-3) that are expressed during development, with high expression in neuronal cells^9,10^. Though there is no current functional validation of IGF2BPs to specific neuropathologies, *IGF2BP3* is a primary microcephaly candidate^11^, and all IGF2BPs are associated with multiple targets connected to autism spectrum disorder^12^. There is also much model organism evidence to support critical roles *IGF2BP*s in brain development. *IGF2BP1* influences pluripotent stem cell adhesion and survival^9,10^, neural stem cell maintenance^13^, hippocampal dendrite arborization^14^, and axon guidance and outgrowth^15,16^. IGF2BP2 localizes in dendrites^17^ and in neural precursors, where it mediates neurogenic and astrocyte differentiation potential^18^. The single ortholog in *Drosophila* (*Imp*) is structurally conserved with vertebrate *IGF2BP*s^19^ and is a well-documented temporal factor in neural stem cells, where it is expressed in a high to low gradient starting in embryogenesis^20^. Descending Imp levels through the larval stage defines stem cell growth and proliferation ^21,22^, neuronal fate and diversity^21,23,24^, and stem cell decommissioning^21,22^. Specifically, *Imp* promotes *Chinmo* translation and consequently represses *mamo* transcription^25^ to control the specification of early and late-born neurons^25^. Imp also controls the growth and proliferation of neural stem cells by stabilizing *Myc* transcripts^22^ and localizes to axons suspected to regulate regrowth during axon remodeling via *prolifin* mRNA transport and localization^26^. In both stem cells and intermediate neural progenitors, *Imp* facilitates proper neuropil targeting to the major learning and memory centers in the adult brain^27^ and is required for proper development of adult olfactory navigation circuitry^28^. Together, these studies show that *Imp* function in neural stem cells during development affects both neuronal morphology^26,29^ and neuronal fate^25,27,30^ in the adult.

While previous work highlights the critical role that *IGF2BP*s/*Imp* have on neurodevelopment, research to date has primarily focused on neural stem cells or intermediate neural progenitors. Furthermore, the functional consequences of developmental *Imp* loss are vastly understudied. This leads us to ask: how does *Imp* regulate neurons both developmentally and functionally, and how does disruption of *Imp* in these neurons correlate to human neuropathologies? We hypothesize that *Imp* is required in neurons themselves for normal neuronal function and to test this hypothesis, we leveraged the *Drosophila* model system to spatially and temporally knock down *Imp* expression. We found that neuronal *Imp* is required during development to ensure proper neuronal function that is specific to seizure behavior. We also found a smaller role for Imp independent of development, because the loss of *Imp* in adulthood caused seizures. We also identified that loss of Imp target, *Syndecan* (*Sdc)*, caused seizures and showed with rescue assays that *Sdc* functionally interacts with *Imp* in seizure behavior. Together, we show that neuronal *Imp* is essential in developing neurons for adult neuronal function and that the interaction between *Imp* and *Sdc* modulates seizure behavior.

## RESULTS

### Loss of Imp causes seizures with normal gross motor function

*Imp* has been highly characterized in neural stem cells where it functions as an important temporal factor to influence the type of neurons produced at different developmental time points^25^. However, it is unknown how loss of neuronal *Imp* might affect brain function independent from its role in stem cells. To test *Imp’s* neuronal function and resulting behavior outputs, we assessed seizure phenotypes using vortex assays. We reduced *Imp* using *in vivo* RNA interference (RNAi) in two primary cell types: neural stem cells and post-mitotic neurons. While loss of *Imp* in both cell types resulted in seizures (Figure 1A, Kruskal-Wallis, *P*= 0.017 and *P*<0.0001 for recovery time compared to eGFP RNAi controls in NSCs and neurons, respectively), loss in post-mitotic neurons caused both a higher frequency of seizures and a trend of longer recovery from seizures (Figure 1B). The average length of seizure was 14.83 seconds in animals with *Imp* knockdown in neural stem cells. However, *Imp* knockdown in neurons caused seizures for an average of 21.77 seconds, correlating neuronal *Imp* loss to a more severe phenotype^31^. The difference in recovery time between stem cell and neuronal *Imp* knockdown was not significant when all individuals were included (Kruskal-Wallis, *P*>0.0009, Figure 1A), but if non-seizing individuals were excluded, post-mitotic neuronal knockdown animals took significantly longer to recover than control knockdown animals in the vortex assay (Mann-Whitney U, *P*=0.0478). Most neural stem cell knockdown animals recovered in under ten seconds and did not differ from the control group. Their seizure episodes were very short, and once animals recovered, they did not experience additional seizures (see supplemental videos 1-2). Neuronal knockdown animals had a more conspicuous phenotype. Noticeable “tonic-clonic” phases with paralysis were interrupted by multiple bouts of spasms during the recovery phase^32^. Individuals experiencing multiple episodes demonstrated periods of flips, brief walks, and a return to supine position, which corresponded to “clonus-like” activity (see supplemental videos 3-4). Neuronal knockdown produced a more robust and consistent seizure phenotype, supporting the idea that neuronal Imp regulates neuron function in the context of seizure behavior. Furthermore, this function was not dependent on *Imp*’s role in neural stem cells. As a result, the remainder of the analyses were performed only on neuronal knockdown animals.

**Figure 1.**
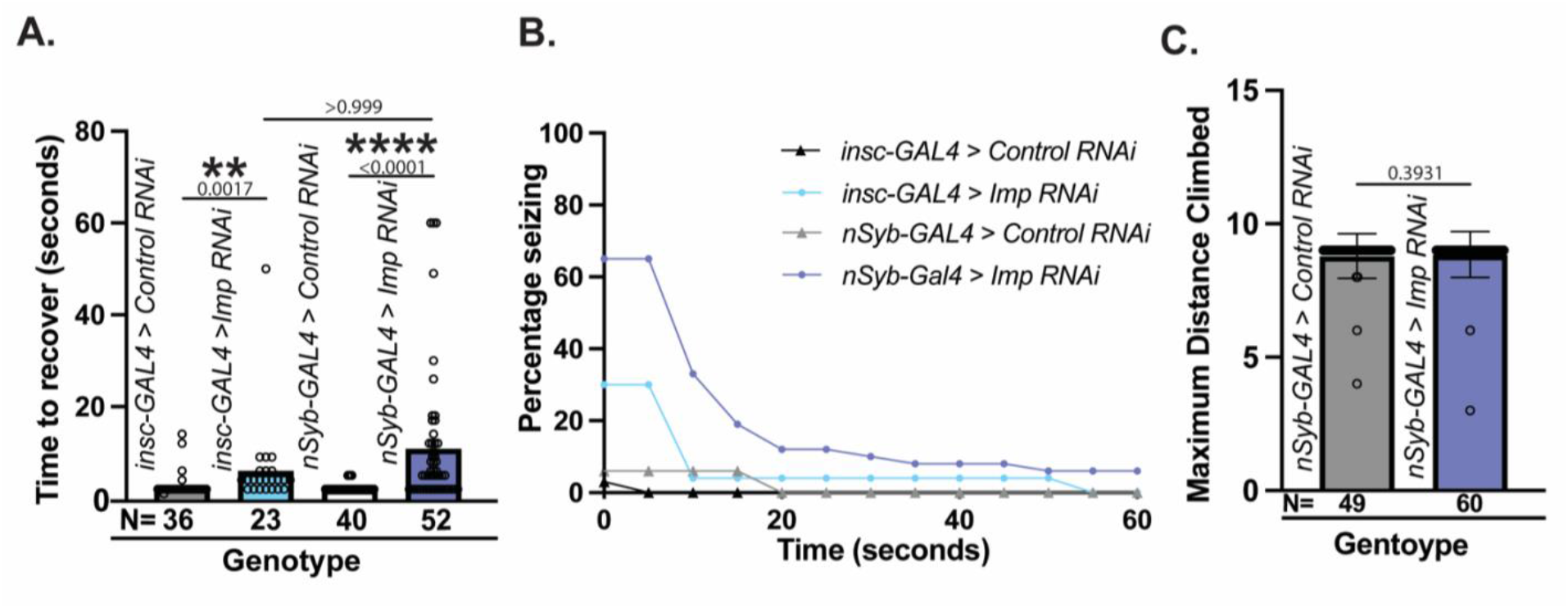
*Imp* knockdown causes higher occurrence and longer duration of seizures with normal gross motor function. (A) Seizure behavior reported as average time to recover after vortexing for *eGFP* RNAi control (*VALIUM22*-*EGFP*.*shRNAI*.*1*) and *Imp* RNAi (*TRIP*.*HMC03794*) expressed using neural stem cell driver *inscuteable*-*GAL4* (*insc-GAL4*) or pan-neuronal driver *neuronal synaptobrevin-GAL4* (*nSyb-GAL4*). Individual points represent each fly, whiskers represent 95% confidence intervals, and bar heights equal the mean. Kruskal-Wallis test determined significance between all conditions. Relevant comparisons reported (p<0.05, ***p<0.01, ***p<0.001,****p<0.0001, ns=p>0.05). N= number of total animals. (B) Percentage of flies seizing reported in (A) observed at 5-second intervals across the entire 60-second trial. Ns are the same as in (A). Seizing is defined by supine paralysis, spastic movement, and/or inability to walk with excessive tremoring. Behavioral assays were performed between days 12 and 14. (C) Maximum distance climbed (cm) in 30 seconds for *eGFP* RNAi control and *Imp* RNAi expressed in *nSyb-GAL4*. Individual points represent individual flies, whiskers represent 95% confidence intervals, and bar heights equal the mean. P-values are reported above each bar. Mann-Whitney U determined significance (p<0.05, ***p<0.01, ***p<0.001,****p<0.0001, ns=p>0.05). N= number of total animals.

Flies have a natural tendency to climb that is inhibited if animals are experiencing gross motor defects^33^. Climbing deficiencies in *Imp* knockdown animals could suggest a general motor deficit rather than a specific functional defect. To differentiate between gross deficits and acute, induced functional disruptions, we used forced climbing assays. Aged flies were placed into vials, banged to the bottom, and allowed to climb to a maximum distance (9mm) for 60 seconds to determine if animals had any broad motor deficits that prevented climbing. We found that loss of neuronal *Imp* did not affect climbing ability (Mann-Whitney U, *P*=0.3931, Figure 1C), suggesting that the observed mechanically induced seizure phenotypes were not due to gross motor problems. Therefore, neuronal Imp functions in seizures but not general motor function.

### Developmental Imp loss is the primary contributor to seizure behavior

RNA binding proteins are broadly acting genes affecting both cell development and mature cell function^5,6,34^. To determine when *Imp* loss results in seizures, we used the GAL4-UAS system to temporally remove *Imp* in neurons. Some *GAL4-UAS* combinations have a temperature-sensitive effect where strong activity occurs at high temperatures (high activity, 29°C) and little-to-no activity occurs at low temperatures (low activity, 18°C). We verified temperature sensitivity by comparing recovery times in vortex assays between control and *Imp* knockdown animals at high activity and low activity conditions. We found a significantly longer recovery time using the high activity conditions (29°C, Kruskal-Wallis *P*<0.0001) but not the low activity conditions (18°C, *P*=0.0554), showing that *nSyb-GAL4* combined with our *Imp* RNAi construct has a temperature sensitive phenotype that can be used to modulate the severity of *Imp* knockdown (Figure 2A). We used this temperature-sensitive phenotype to disentangle *Imp* function during development vs in the adult. We placed developing animals in the low activity condition (18°C) and moved adults to the high activity condition (29°C) upon eclosion. Our schedule was designed to isolate *Imp* knockdown to adulthood only. Seizure activity increased in animals with *Imp* knockdown only during adulthood, but it was not significantly different from its eGFP RNAi control match (Kruskal-Wallis, *P*=0.0813). Next, we placed developing animals in the high activity condition (29°C) during development and upon eclosion moved adults to the low activity condition (18°C). This time, our schedule was designed to isolate *Imp* knockdown to development only. Developmental *Imp* knockdown caused significantly longer recovery times when compared to eGFP RNAi knockdown controls (Kruskal-Wallis *P*<0.0001). Interestingly, developmental *Imp* and adult *Imp* knockdown group recovery times were not significantly different from each other even though the average developmental *Imp* recovery time (21.06 seconds) was approximately twice as long as adult *Imp* knockdown (10.46 seconds, Kruskall-Wallis *P*>0.999). However, there was a much higher proportion of individuals seizing in the developmental knockdown group compared to the adult knockdown group (Figure 2B). Our results suggest that *Imp* functions to u seizures during both development and in the adult, although to an apparent lesser degree in adults. In addition, developmental *Imp* knockdown (29°C >18°C) more closely phenocopies continual knockdown (29°C >29°C) phenotypes. Together, these results suggest that developmental Imp expression plays a critical role in programming future neural function, and both developmental and adult Imp expression could contribute to neuronal function important for seizure regulation.

**Figure 2.**
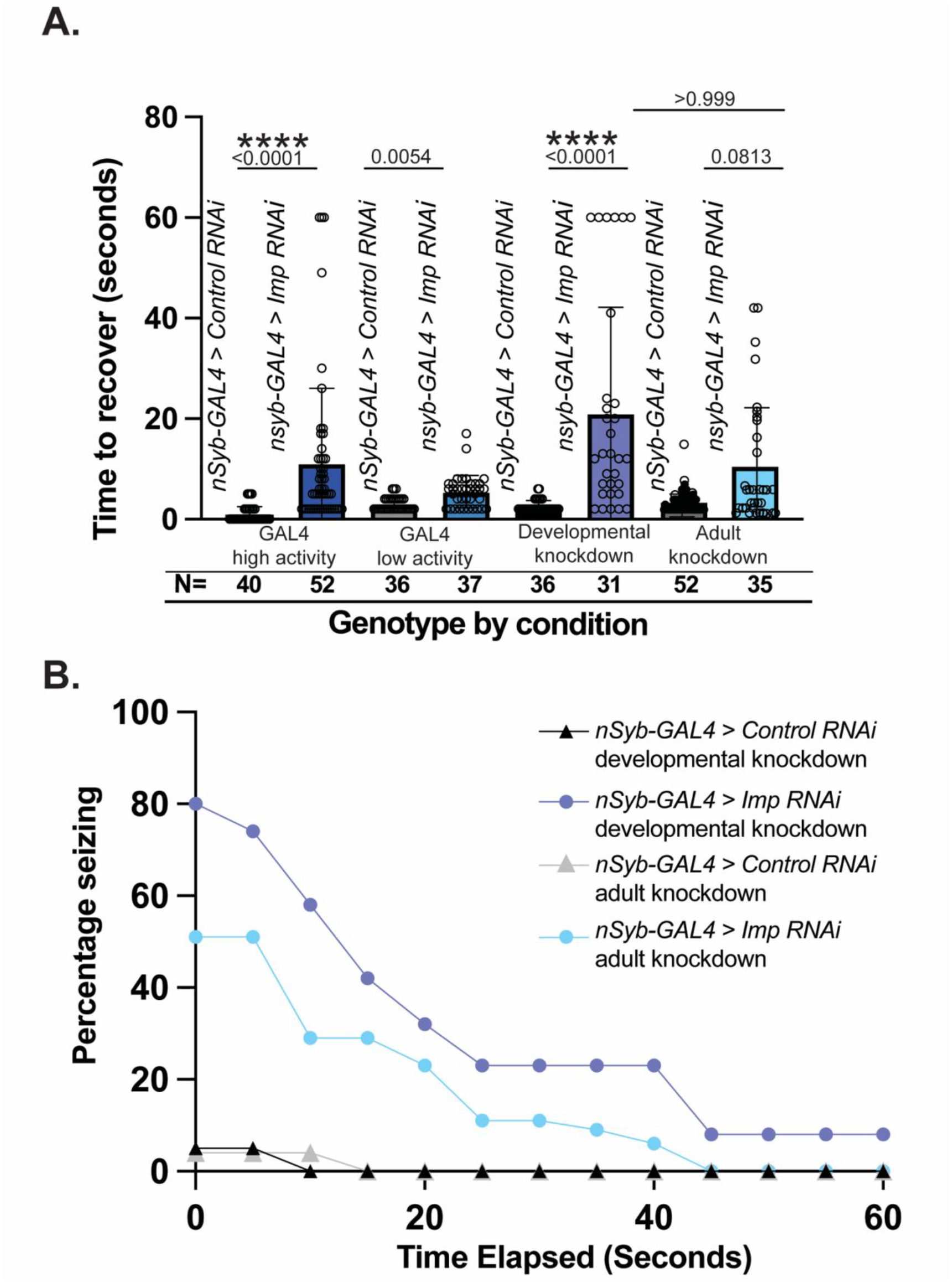
Developmental loss of neuronal Imp is the primary source of seizure behavior. (A) Average time to recover for flies exhibiting seizure behavior for neuronal (*nSyb-GAL4*) *eGFP* RNAi (control) and neuronal *Imp* RNAi. High GAL4 activity animals are raised at 29°C and low GAL4 activity animals are raised at 18°C. Results demonstrate that *nSyb-GAL4* RNAi is temperature sensitive. Developmental knockdown animals are raised at 29°C and moved to 18°C within one day of adulthood. Adult knockdown animals developed at 18°C and shifted to 29°C within four hours of adulthood. Behavioral assays were performed at day 14. Individual points represent individual flies, whiskers represent 95% confidence intervals, and bar heights equal the mean.. P-values are reported above each bar. Kruskal-Wallis determined significance between all conditions. N= number of total animals. Relevant comparisons reported (p<0.05, ***p<0.01, ***p<0.001,****p<0.0001, ns=p>0.05). Developmental knockdown of *Imp* is sufficient to recapitulate the seizure phenotype. (B) Percentage of flies presented in (A) seizing observed at 5-second intervals across the entire 60-second trial.

### The Imp downstream target Sdc is required for normal neuronal function and interacts with Imp molecularly and functionally

Since Imp is an RNA-binding protein, Imp’s downstream target genes must be identified to molecularly understand Imp’s control of seizures. Samuels *et al*. (2020)^35^ isolated mRNAs that bound to the Imp protein in third-instar larval brains. We leveraged this list of potential Imp mRNA targets to identify relevant candidates in seizure behavior using RNAi and vortex assays. *Sdc* knockdown was particularly interesting because it had a significantly longer recovery time compared to controls (Figure 3A, Mann-Whitney U *P*<0.0001) and exhibited robust tonic-clonic episodes after vortexing (see supplemental video 5), suggesting that *Sdc* also is required to properly regulate seizure behavior.

**Figure 3.**
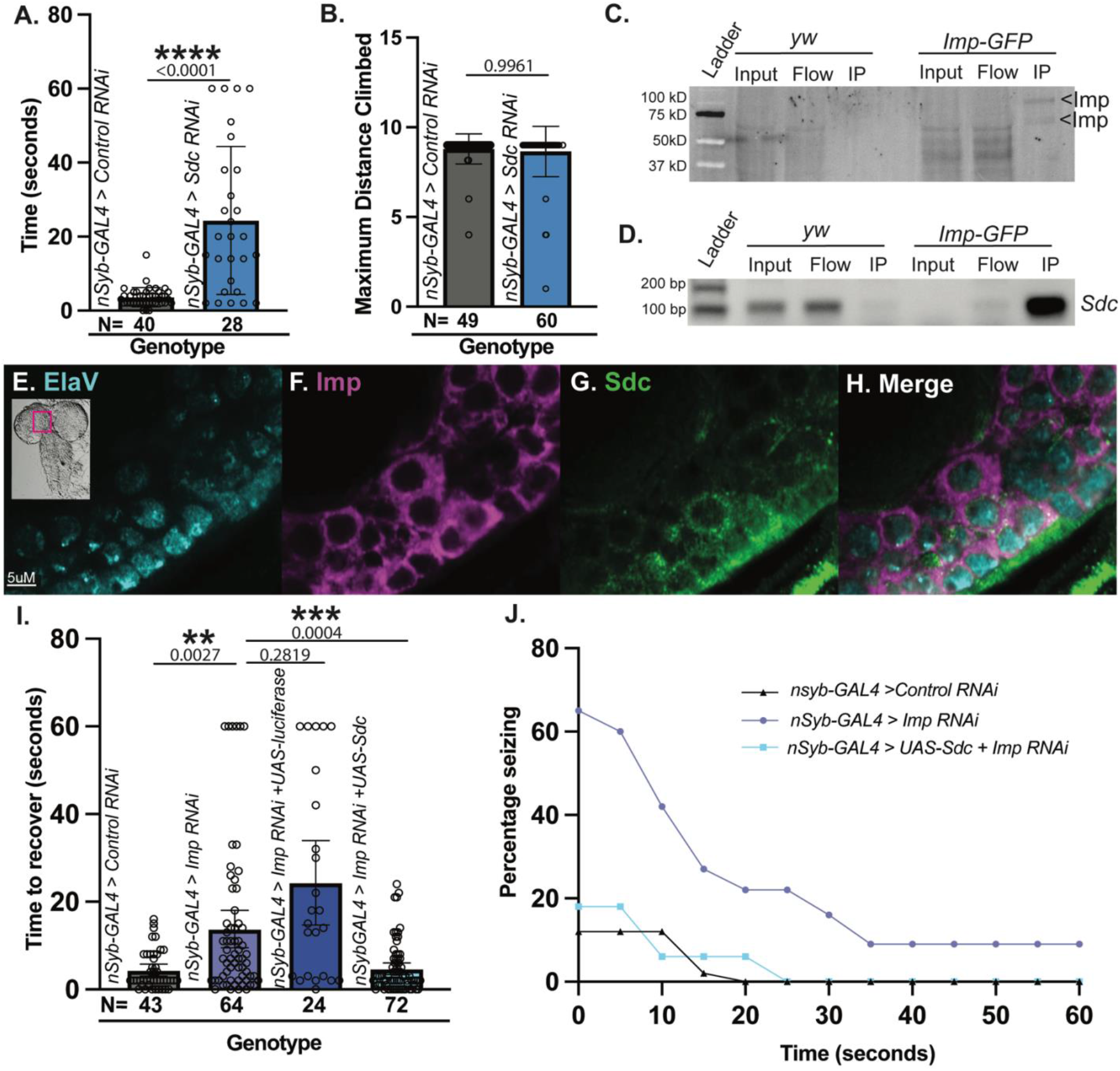
Sdc interacts with Imp molecularly and functionally and is required for normal neuronal function. (A) Average time to recover for flies exhibiting seizure behavior for neuronal (*nSyb-GAL4*) *eGFP* RNAi (control) and neuronal *Sdc* RNAi. Individual points represent each fly and whiskers represent 95% confidence intervals. N= number of total animals. Mann-Whitney U determined significance (p<0.05, ***p<0.01, ***p<0.001,****p<0.0001, ns=p>0.05). (B) Maximum distance climbed (cm) in 30 seconds for *eGFP* RNAi control and *Sdc* RNAi. Mann-Whitney U significance (*p<0.05, ***p<0.01, ***p<0.001,****p<0.0001, ns=p>0.05). (C) RNA immunoprecipitation of Imp-GFP protein from control (*yw*) and *Imp-GFP* animals using nanobody GFP agarose. Imp is immunoprecipitated specifically in *Imp-GFP* animals. (D) PCR analysis of Imp-GFP RNA immunoprecipitation using primers specific to *Sdc* in control (*yw*) and *Imp-GFP* animals. *Sdc* product is amplified in *Imp-GFP* RIP but not in control RIP (*yw*). (E-H) Confocal images of a single slice through the central brain of a third instar larva stained with ElaV (neurons, cyan), Imp (magenta), and Sdc-GFP (green). Imp and Sdc proteins are both present in neurons. (I) Average time to recover after vortexing for flies exhibiting seizure behavior for neuronal expression (*nSyb-GAL4*) for *eGFP* RNAi (control), *Imp* RNAi, *Imp* RNAi + *UAS-Luciferase* (dilution effect control), and *Imp* RNAi + *UAS-Sdc* cDNA. Points represent individual flies, whiskers represent 95% confidence intervals, and bar heights equal the mean. P-values are reported above each bar. N= number of total animals. Kruskal-Wallis determined significance between all conditions, relevant comparisons reported (p<0.05, ***p<0.01, ***p<0.001,****p<0.0001, ns=p>0.05). (J) Percentage of flies seizing presented in (E) observed at 5-second intervals across the entire 60-second trial. Expression of Sdc cDNA in Imp knockdown animals rescues seizure deficits.

*Sdc* encodes a heparan sulfate proteoglycan that regulates midline axon guidance during development^36^ and the development of the neuromuscular junction^37^, indicating that it could also function in gross motor behavior. However, the strongest candidate for downstream target analysis would cause seizure-specific deficiencies, but not gross motor deficits. To ensure we chose a strong candidate and rule-out broad motor defects, we performed forced climbing assays on *Sdc* neuronal knockdown animals. Like *Imp, Sdc* knockdown did not cause any gross motor defects (average recovery=24.76 seconds, Figure 3B, Mann-Whitney U P=0.9961). Therefore, *Sdc* is a strong candidate for seizure regulation. If *Sdc* mRNA is a target of Imp, Imp protein should bind *Sdc* mRNA. We performed RNA Immunoprecipitation (RIP) in third-instar larval brains using Imp-GFP animals and anti-GFP agarose to validate that Imp binds to *Sdc* mRNA. Western analysis of RIP samples show Imp-GFP is sufficiently immunoprecipitated (Figure 3C). In addition, we found signal for *Sdc* in Imp-GFP pulldowns, but not when using a control fly line that does not contain GFP (*yw*, Figure 3D). These data suggest a strong and specific physical interaction between Imp and *Sdc* mRNA. Next, we identified whether Imp and Sdc were co-expressed in neurons in the brain. We immunostained third-instar larval brains for Imp, Sdc, and Elav (to mark neurons). We found that Imp and Sdc co-express in central brain neurons (Figure 3E-H). Co-expression of proteins in neurons supports a relevant functional interaction, and our data indicate that together, Imp and Sdc regulate neuronal function.

Physical interaction between Imp protein and *Sdc* mRNA combined with co-expression in relevant cell types is highly suggestive of a functional relationship. However, to conclude that the observed interaction is biologically relevant for our phenotype of interest, we tested whether Sdc overexpression could rescue *Imp*-induced seizures. We quantified phenotypic rescue in flies expressing *Sdc* cDNA while simultaneously knocking down *Imp* in neurons to identify such relevant interactions. *Sdc* cDNA expression significantly reduced the time needed to recover from vortexing in *Imp* knockdown animals (average time to recover=4.722 seconds, Figure 3I) compared to *Imp* knockdown alone (average time to recover=13.77 seconds, Kruskal-Wallis *P*=0.0004). In addition, the percentage of flies seizing was reduced to near control levels (Figure 3J) and tonic-clonic episodes were nearly absent (see supplemental video 6). Our data indicate complete rescue of seizure deficits and strongly suggest that Imp regulates *Sdc* mRNA in the context of neuronal function and seizure behavior.

## DISCUSSION

Identifying genes that contribute to seizure disorders is an important first step in understanding how they arise and can best be treated. In this study, we demonstrate that Imp and its downstream target *Sdc* are required in neurons for seizure behavior regulation. Our study is one of few studies that has linked developmental Imp to adult behaviors^38^ and highlights the critical role of both Imp and Sdc in post-mitotic neuronal development. We can now tease apart the critical developmental pathways through which Imp regulates proper neuronal function.

To identify the nature of how seizures are caused, it is important to investigate the contribution of individual cell types during development. *Imp* function in neural stem cells is consequential to many neurodevelopmental outcomes, including seizure phenotypes. Interestingly, our results suggest that *Imp* expression in post-mitotic neurons is most critical to prevent seizures; loss of *Imp* in neural stem cells only has a small effect on adult neuronal function. Because little is known about the role of Imp in neurons, how *Imp* regulates development and function in the context of the terminally fated cell is an open question. We can, however, gain insight from our findings. Imp binds *Sdc* mRNA, and *Sdc* phenocopies *Imp* seizure phenotypes. These data suggest that *Imp* disruption directly impacts *Sdc*-specific phenotypes to cause seizures. Sdc is a conserved transmembrane heparan sulfate proteoglycan^39^ that is secreted from midline cells of the fly central nervous system and binds products that mediate axon guidance^37^. *Sdc* also promotes synapse growth at the neuromuscular junction^36^. Given that loss of both *Imp* and *Sdc* cause seizure phenotypes, we posit that Imp expression is required for proper neuronal growth because it regulates *Sdc* expression. Modifications in neural structure, particularly dendritic length and/or branching pattern, can change its firing pattern^40^. A vertebrate paralog of *Sdc* (Syndecan-2) promotes the formation of dendritic spines, and so would be interesting to test Syndecan-2’s role in neuronal activity and seizure phenotypes. *Sdc*’s role at the neuromuscular junction is post-synaptic^37^, so it is possible that *Imp* influences dendrite growth required for proper neuronal function via *Sdc* expression levels. Alternatively, loss of *Imp* and/or *Sdc* in post-mitotic neurons could disrupt cell survival and induce loss of neurons. Neuronal loss has been linked to asynchronization and tonic depolarization leading to seizures^41^. Whether loss of *Imp* causes supernumerary neuron cell death that leads to seizures, and whether *Sdc* overexpression could rescue this phenotype is intriguing, and an essential future study.

Though *Imp* has been studied extensively in a developmental context, our data suggest that *Imp* plays a role in adult neuronal maintenance and function. This study sought to determine when *Imp* function was required for seizure regulation. Our results strongly suggest that developmental *Imp* has the most influence on neuronal function later in life. However, flies with reduced Imp only in adulthood still seized while loss of developmental *Imp* caused more severe seizures with increased duration and a higher likelihood of occurrence. Our results are in line with what is known about *Imp* being a key factor in early brain development that influences adult brain circuitry^27^ and behavior^38^. Beyond development, RNA-binding proteins play roles in cell migration, maturation, and synaptic integration in mature brain cells^34^. *Imp*, therefore, could be a key player in neuronal maintenance that suppresses seizures. Identifying changes in neuronal circuitry as well as excitation and inhibition imbalances in animals with only adult Imp loss will be a first important step in disentangling the functional role of *Imp*.

The seizures observed in both *Imp* and *Sdc* neuronal knockdown animals were relatively short but conspicuous and stereotyped. Clear tonic-clonic episodes were observed in all trials. Previous *Drosophila* epilepsy studies suggest that time to recover determines the severity of seizures ^2^. However, seizure length is a superficial way to characterize seizure phenotype. Additional details provide a more comprehensive view of *Imp*-related seizures. For example, more severe bang-sensitive mutants such as *bss* take over 500 seconds to recover from the same mechanical perturbation but also experience gross climbing defects ^42^, making it difficult to assess the severity of seizure independent of gross motor phenotype. *Imp* knockdown animals, on the other hand, do not show motor deficits, so we can leverage this seizure-specific phenotype to probe mechanisms specific to seizure activity.

We provided evidence for a functionally relevant interaction between *Imp* and *Sdc* in seizure regulation, but we have yet to identify the mechanism by which *Imp* mediates Sdc expression. RNA-binding proteins modulate gene expression by stabilizing downstream mRNAs, modifying transcription and translation, and transporting mRNAs to different parts of the cell^5,6^. Multiple studies have identified that *Imp* controls neurodevelopmental processes by regulating downstream target mRNA expression^21,22,25,30,30,43^ but the mechanisms regarding *Sdc* mRNA have not been explored. Because *Sdc* cDNA overexpression rescues seizure phenotypes in *Imp* knockdown, we hypothesize that *Imp* positively regulates Sdc protein levels. Molecular assessment of how loss of Imp changes *Sdc* mRNA and protein levels, as well as any changes in *Sdc* mRNA localization, will elucidate the mechanism by which Imp regulates *Sdc* during development to control seizure behavior.

Like many RNA-binding proteins, *Imp* has numerous downstream targets. In addition to *Sdc*, some Imp targets such as *Para*^3,44–47^ and *Shaker*^48–50^ have previously been characterized for their role in seizure biology. Samuels et. al (2020) identified an additional 37 Imp targets in larval brains, so other pathways could be involved in seizure control. Additionally, other cell types could affect seizure regulation. Glia, for example, provide critical support for extra-cellular matrix (ECM) remodeling and perineuronal net integrity^51–54^ both which influence seizure occurrence^55^. Given that heparan sulfate proteoglycans such as *Sdc* are ECM proteins ^39,56^, it will be important to investigate the role of both *Imp* and *Sdc* in glia during development and in adult brain function.

In conclusion, we show a role for the RNA-binding protein *Imp* in post-mitotic neuronal regulation of brain function, where loss of neuronal Imp expression causes seizure behaviors. Additionally, we provide support for a functionally relevant interaction between *Imp* and *Sdc* in seizure behavior. Our work contributes to a growing body of literature highlighting the role of RNA binding proteins as critical regulators of brain function and opens many new exciting questions about how *Imp* and other RNA binding proteins could regulate brain development and function through its role in post mitotic neurons.

## ACKNOWLEDGMENTS

We thank members of the Link lab for critical reading of the manuscript. P.R. was supported by the Eunice Kennedy Shriver National Institute of Child Health & Human Development of the National Institutes of Health under Award Number (T32HD007491) and the National Institutes of Health under Ruth L. Kirschstein National Research Service Award T32HG008962 from the National Human Genome Research Institute at the University of Utah. The content is solely the responsibility of the authors and does not necessarily represent the official views of the National Institutes of Health. N.L. was funded in-part by NIAID grant R56AI170857 and R01AI170857 and the University of Utah. We thank Claude Desplan for providing the anti-Imp antibody used in this study. The monoclonal antibody Elav-9F8A9, developed by Gerald M. Rubin at Janelia Farm Research/HHMI was obtained from the Developmental Studies Hybridoma Bank, created by the NICHD of the NIH and maintained at The University of Iowa, Department of Biology, Iowa City, IA 52242. We acknowledge the information provided by FlyBase (NHGRI awards U41HG000739 and U24HG010859) using release FB2023_04. Stocks obtained from the Bloomington Drosophila Stock Center (NIH P40OD018537) were used in this study.

## METHODS

### Drosophila melanogaster strains and culture

The following lines were used in this study with genotypes separated by commas: *w[*]*; *P{w[+mW*.*hs]=GawB}insc[Mz1407]* (RRID: BDSC_8751), *y[1] w[1118]; P{y[+t7*.*7] w[+mC]=nSyb-GAL4*.*P}attP2* (RRID: BDSC_51941), *y[1]sc[*] v[1] sev [21]; P{y[+t7*.*7] v[+t1*.*8]=VALIUM22-EGFP*.*RNAi*.*1}attP40* (RRID: BDSC_41557), *y[1]sc[*] v[1] sev [21]; P{y[+t7*.*7] v[+t1*.*8]=TRIP*.*HMC03794}attP40* (RRID: BDSC_55645), *P{KK108799}VIE-260B* (RRID:Flybase_FBst0479142), *w[*]; P{w[+mC]=UAS-Sdc*.*J}3* (RRID: BDSC_8564), *P{UAS-t, y[1]w[*] Mi{PT-GFSTF*.*2}Imp{M105901-GFSTF*.*2* (RRID: BDSC_60237), *y[1] w[*]; Mi{PT-GFSTF*.*0}Sdc[MI10787-GFSTF*.*0]/Cyo (RRID: BDSC_66373)*.

All lines were maintained at either 25°C or 18°C on Archon glucose medium supplemented with dry yeast in wide plastic vials. All incubators were kept on a 12:12 hour light:dark cycle.

### Behavior Assays

#### Standard behavior conditions

All flies were aged between 12 and 14 days (see each assay method for details), where all flies were age-matched between control and knockdown groups for each trial set. Carbon dioxide can acutely alter behavior^57^, so flies were only anesthetized with carbon dioxide when collected as virgins and later moved into test vials using mouth pipetting. All behavior trials were performed within 4 hours of lights on. Females exhibited a more consistent phenotype in *Imp* pilot seizure assays (see figure S1), so only females were used for data collection. All trials were recorded using either an iPhone 13 mini or a Panasonic DMC-G85 camera with an Olympus ED 60mm f2.8 macro lens.

#### Seizure Assays

Female progeny from each experimental genotype were matched to the same driver control knockdown (*eGFP*) and aged to 12 days at 29°C. Standard bang sensitivity assays, specifically vortex assays, were performed to assess seizure behavior^2^ where vials were vortexed for 10 seconds and placed on the bench at eye level to easily record flies. The proportion of flies seizing and time to recovery were recorded for each vial. Seizing was defined by supine paralysis, spastic movement, and/or inability to walk with excessive tremoring. Recovery was defined as walking normally in the correct orientation without abdominal or wing spasms and/or exhibiting normal geotaxis behavior without abdominal/wing spasms or falling. Some flies climbed very quickly after flipping over and did not walk along the bottom of the vial. Trials were ended once all flies had recovered fully or until 60 seconds.

#### Developmental/Functional Assays

*Imp* RNAi crosses were set, and progeny developed at either 18°C (low GAL4 activity temperature) or 29°C (high GAL4 activity temperature). Females were separated, placed in individual vials within four hours of eclosion, and either left at their developmental temperature or moved to the inverse temperature to age for 12-14 days. All trial sets were age matched between control and RNAi genotypes. On the assay day, aged flies were aspirated into empty vials in sets of up to 10 and acclimated for 20 minutes at their aged temperature (18°C or 29°C). Flies were analyzed for seizure using the standard bang assay described in the initial knockdown assays.

#### Rescue Assays

Some complex genotypes were sick when crossed at 29°C, so crosses were set, and progeny developed at 25°C to ensure optimal animal health while still allowing sufficient GAL4 activity. Female progeny were collected within eight hours of eclosion and aged to 14 days at 25°C. On seizure assay days, aged flies were aspirated into empty vials in sets of up to 10 and acclimated for 20 minutes at 25°C. Flies were analyzed for seizure using the standard bang assay described in the initial knockdown assays.

#### Forced-climbing Assays

Female progeny developed and aged 14 days at 25°C. On the day of testing, aged female flies were aspirated into empty vials marked with 1cm lines in sets of up to 10 and acclimated for 20 minutes at 25°C. Vials were banged on the benchtop 3 times to ensure that all flies were at the vial bottom and were allowed 60 seconds to climb to their maximum distance (up to 9cm). The maximum distance climbed by each individual fly was recorded.

### RNA Immunoprecipitation

40 *yw* (control) and *y[1]w[*] Mi{PT-GFSTF*.*2}Imp{M105901-GFSTF*.*2* (RRID: BDSC_60237) larvae were collected, fixed in 0.1% paraformaldehyde (EM Grade, Electron Microscopy Sciences #15710) in 1X Phosphate Buffered Saline (PBS) + 0.3% Triton X-100 for 10 minutes. From here, all steps were carried out in an RNAse free manner with RNAse inhibitors if needed. Larvae were rinsed with 0.125M Glycine to stop fixation, rinsed in PBS, and transferred to Chaps cell extract buffer (Cell Signaling Technology #9852) with Pierce phosphatase and protease inhibitors (Thermo Scientific #A32961) or Halt protease and phosphatase inhibitor cocktail (Thermo Scientific #78440). Animals were ground with a motorized pestle, frozen at -80°C, thawed on ice, and vortexed three times. Debris was pelleted by centrifuging for 10 minutes at 4°C.

Supernatant was transferred to a new tube. 50μl was saved for RNA and protein input samples. Chromotek GFP-Trap Agarose kit was used to immunoprecipitate Imp-GFP and wash sample. 50μl of lysate flow was saved for RNA and protein analysis. 1/3 of immunoprecipitate was saved for protein analysis. To disassociate RNA from Imp-GFP/beads, GFP agarose was resuspended in 100ul Lysis Buffer (from Chromotek kit) with 30μg Proteinase K and incubated for 30 minutes at 55°C in a thermoshaker with agitation. RNA was extracted using Trizol (Thermo Scientific #15596026). 600μl of Trizol was added, mixed by pipetting up and down, and incubated at room temperature for 5 minutes. 150μl of chloroform was added, tubes were shaken vigorously for 15 seconds, and samples incubated for 10 minutes at room temperature. Tubes were centrifuged for 15 min at 4°C and aqueous phase was removed and saved. 1.5μl of GlycoBlue and 500μl of isopropanol were added and tubes were vortexed for 10 seconds. Samples incubated overnight at -20°C and centrifuged for 15 minutes at 4°C for precipitation. RNA pellets were washed with 70% ethanol and dried pellets were resuspended in 20μl of RNAse free water. cDNA was generated using Bio-Rad iScript cDNA synthesis kit (Bio-Rad #1708890). Primers specific for Sdc were used for PCR (Forward: TCCTACGGCCATAGCCATTAT Reverse: CGGTAGAAGTAATTCCGCCAGA)

### Larval Dissection and Immunohistochemistry

Brains from late third instar larvae (identified by gut clearing and spiracle protrusion) were dissected in cold Phosphate Buffered Saline (PBS) and fixed in 4% paraformaldehyde in PBS + 0.3% Triton-X (PBST) for 20 minutes, then washed in PBST for five minutes three times before blocking in PBST + 1% Bovine Serum (PBSTB) for 30 minutes two times. Brains were blocked a third time in PBSTB + 5% Normal Donkey Serum (NDS) for 30 minutes. Brains were incubated in primary antibodies diluted in PBSTB for 3 days at 4°C, washed with PBSTB for 20 minutes three times, and then incubated in secondary for 2 days at 4°C. Brains were washed in PBST four times for 15 minutes, adding DAPI into the second wash, and mounted in Slow Fade Gold on microscope slides with spacers made from a single layer of double-sided tape.

We used the following antibodies in this study: rabbit anti-GFP (1:1000, Invitrogen A-11122, RRID: AB_221569), rat anti-Imp (1:200, gifted from Dr. Claude Desplan, New York University), mouse-anti ElaV (1:100, Developmental Studies Hybridoma Bank 9F8A9, RRID: AB_2314364), Donkey anti-rabbit 488 (1:500, Jackson ImmunoResearch 711-545-152, RRID: AB_2313584), Donkey anti-rat fluorophore 647 (1:500, Jackson ImmunoResearch 712-605-153, RRID: AB_2340694) Donkey anti-mouse fluorophore Rhodamine Red™-X secondary (1:500, Jackson ImmunoResearch 715-295-151, RRID: AB_2340832).

Brains were imaged on a Zeiss LSM980 confocal microscope using Airyscan. Single 2um slices through the middle of the brain were imaged with a 63x oil objective.

### Statistics

For seizure assays, the total time to recovery was analyzed between control and treatment groups using a Mann-Whitney U for single comparisons a Kruskall-Wallis for multiple comparisons. For motor assays, maximum distance climbed was analyzed between control and treatment groups with a Mann-Whitney U single comparison test. We chose non-parametric tests because the variation in behavior within genotypes follow non-normal distributions. All statistical analyses were performed using GraphPad Prism version 10.0 for Mac OS, Graphpad Software, Boston Massachusetts USA, www.graphpadprism.com. Power of test was determined using post-hoc analysis in G-Power^58^ using an Alpha=0.5 and Power set to .80. Numbers for individual experiments corresponding to each figure are reported in Table S1.

