## Supplementary figures and images for "Loss of neuronal Imp induces seizure behavior through Syndecan function"

### Supplemental Figure 1

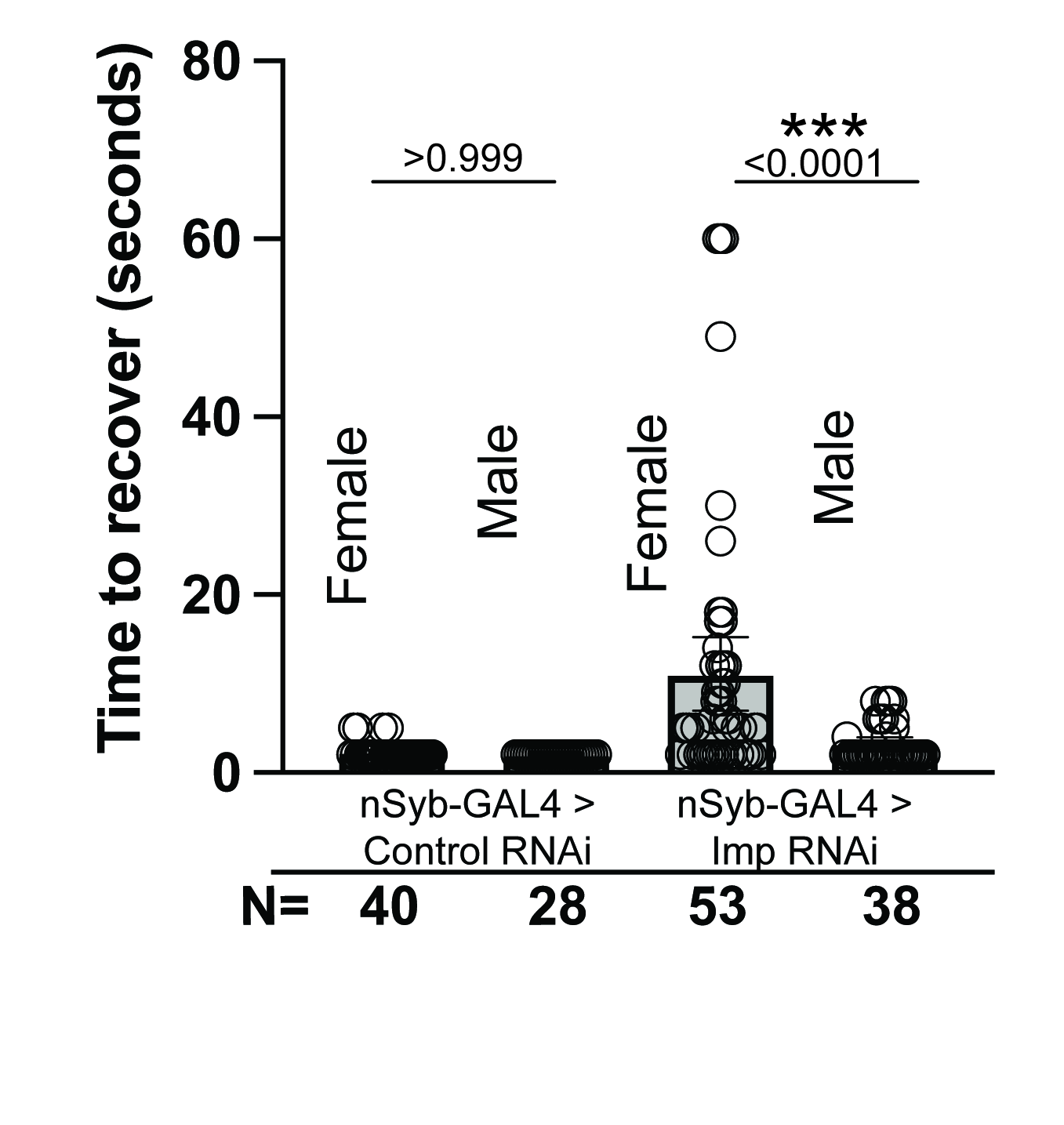
