## Supplemental Figure Captions and Table 1 for "Loss of neuronal Imp induces seizure behavior through Syndecan function"

Supplemental Videos

SV1. Video showing normal baseline recovery from vortexing in neural stem cell knockdown in control animals (eGFP RNAi).
SV2. Video showing slight impairment but relatively quick recovery from vortexing for animals when *Imp* is knocked down in neural stem cells.
SV3. Video showing normal baseline recovery from vortexing in pan-neuronal knockdown in control animals (eGFP RNAi).
SV4. Video showing impairment in recovery from vortexing for animals with *Imp* knocked down in pan-neuronally, with visible tonic-clonic episodes.
SV5. Video showing impairment in recovery from vortexing for animals with *Sdc* knocked down in pan-neuronally, with visible tonic-clonic episodes.
SV6. Video showing recovery from vortexing in animals with *Sdc* cDNA expression in *Imp* pan-neuronal knockdown background, with fewer seizing animals and relatively quick recovery.

Supplemental Figures

Figure S1. *Imp* knockdown has a stronger effect on females. Seizure behavior reported as average time to recover after vortexing for *eGFP* RNAi (VALIUM22-*EGFP.shRNAI.1*) Control and *Imp* RNAi (*TRIP.HMC03794*) expressed using pan-neuronal driver *neuronal synaptobrevin-GAL4* (*nSyb-GAL4*). Individual points represent each fly, whiskers represent 95% confidence intervals, and bar heights equal the mean. Kruskal-Wallis test determined significance between all conditions. Relevant comparisons reported (p<0.05, ***p<0.01, ***p<0.001,****p<0.0001, ns=p>0.05). N= number of total animals.

Supplemental Tables

Table 1. Power analysis results for all behavioral assays determined post-hoc in G-Power

| Experiment | Corresponding Figure | Statistical Test | Power |
|---|---|---|---|
| Bang assay cell specific *Imp* knockdown | 1A | Kruskal-Wallis | 0.9615773 |
| Forced-climbing assay *Imp* knockdown | 1C | Mann-Whitney U | 0.9830928 |
| Bang assay staging *Imp* knockdown | 2A | Kruskal-Wallis | 0.9832125 |
| Bang assay *Sdc* knockdown | 3A | Mann-Whitney U | 0.9999438 |
| Forced-climbing assay Sdc knockdown | 3B | Mann-Whitney U | 0.8732388 |
| Bang assay Sdc rescue | 3I | Kruskal-Wallis | 0.9610394 |
| Bang assay by sex | S1 | Kruskal-Wallis | 0.9998712 |
